# Unique and redundant roles of SOX2 and SOX17 in regulating the germ cell tumour fate

**DOI:** 10.1101/643114

**Authors:** Sina Jostes, Martin Fellermeyer, Gina Merges, Glen Kristiansen, Daniel Nettersheim, Hubert Schorle

## Abstract

Embryonal carcinomas (ECs) and seminomas are testicular germ cell tumours. ECs display expression of SOX2, while seminomas display expression of SOX17. In somatic differentiation, SOX17 drives endodermal cell fate. However, seminomas lack expression of endoderm markers, but show features of pluripotency. Here, we use ChIP-sequencing to report and compare the binding pattern of SOX17 in seminoma-like TCam-2 cells to SOX2 in EC-like 2102EP cells and SOX17 in somatic cells. In seminoma-like cells, SOX17 was detected at canonical (SOX2/OCT4), compressed (SOX17/OCT4) and other SOX family member motifs. SOX17 directly regulates *TFAP2C*, *PRDM1* and *PRDM14*, thereby maintaining latent pluripotency and supressing somatic differentiation. In contrast, in somatic cells canonical motifs are not bound by SOX17. In sum only 11% of SOX17 binding sites overlap in seminoma-like and somatic cells. This illustrates that binding site choice is highly dynamic and cell type specific. Deletion of SOX17 in seminoma-like cells resulted in loss of pluripotency, marked by a reduction of OCT4 protein level and loss of alkaline phosphatase activity. Further, we found that in EC-like cells SOX2 regulates pluripotency-associated genes, predominantly by partnering with OCT4. In conclusion, SOX17 (in seminomas) functionally replaces SOX2 (in ECs) to maintain expression of the pluripotency cluster.

## INTRODUCTION

Testicular cancer is the most common malignancy in young men^1^. The most frequent testicular tumours are the type II germ cell tumours (TGCTs). Based on epigenetic profile, morphology and marker expression TGCTs are classified as seminomas or non-seminomas ^1^. Non-seminomas initially present as embryonal carcinomas (ECs), which can further differentiate into mixed non-seminoma ^1^. Seminomas and ECs arise from the same precursor lesion (germ cell neoplasia in situ / GCNIS), which is believed to result from primordial germ cells (PGCs) arrested in development ^2^. Seminomas retain to the most part the genetic and epigenetic signature of PGCs i.e. they express pluripotency (*LIN28, NANOG, POU5F1* (*OCT4*)) and PGC (*PRDM1, TFAP2C, DMRT1, cKIT*, *SOX17*) markers ^3^. The gene expression profile of ECs is highly similar to embryonic stem cells (ESCs), hallmarked by expression of pluripotency markers (*LIN28*, *NANOG*, *POU5F1* (OCT4), *TFAP2C*, *GDF3*, *DNMT3B*, *TDGF1*, *SOX2*) ^3^. Interestingly, seminomas are SOX17+ and SOX2−, while ECs are SOX17− and SOX2+ ^4^.

Both SOX17 and SOX2 are members of the SOX family of transcription factors, which comprises 20 proteins hallmarked by a high mobility group box DNA binding domain ^5^. Both SOX2 and SOX17 are able to partner with OCT4 and, as a consequence, recognize and bind specific binding motifs ^6,7^. In human and mouse ESCs, SOX2/OCT4 bind to canonical motifs on the DNA (*CTTTGT<u>C</u>ATGCAAAT*-like), which are composite SOX (*CATTGTC*-like) and OCT (*ATGCAAAT*-like) motifs. Canonical motifs are located in several enhancers of pluripotency genes ^7,8^. By SOX2/OCT4 mediated transactivation of pluripotency markers, ESCs maintain their undifferentiated, pluripotent status ^7,8^. An increase of SOX17 levels in ESCs leads to a shift of OCT4 from partnering with SOX2 to SOX17, resulting in binding to compressed SOX17/OCT4 motifs ^7,9^. Compressed motifs (*CTTTGTATGCAAAT*-like) are similarly made up from juxtaposition of SOX and OCT binding sites, but lack a central cytosine ^7^. Compressed motifs are predominantly found in regulatory regions of somatic genes ^7,9^. In mouse and human ESCs, binding of SOX17/OCT4 to compressed motifs initiates endodermal differentiation ^7,9^. Although in ESCs the SOX17/OCT4 heterodimer is preferentially bound to the compressed motif, it can also be detected at the canonical motif ^7^.

In human PGCs SOX17 and OCT4 are also highly expressed ^10^. Alike seminomas, human PGCs lack expression of endoderm-associated genes but display a so-called latent pluripotency, i.e. expressing core regulators of pluripotency while being unable to spontaneously differentiate like ECs or ESCs. Three transcription factors, SOX17, PRDM1 and TFAP2C induce the PGC fate in the early embryo and establish and maintain latent pluripotency ^10^. We reasoned that in human PGCs and seminomas, SOX17 acts not as an endoderm-initiation factor but supports latent pluripotency. In line with our hypothesis, it was shown by Irie *et al*. that knockdown of SOX17 in seminoma cells results in downregulation of PGC / seminoma and pluripotency markers ^10^. We further assumed that SOX2 and SOX17 may fulfil similar roles in maintaining expression of pluripotency genes in EC and seminoma cells, respectively. Since it was shown in mice that the SOX17/OCT4 heterodimer is able to bind to the canonical motif, we hypothesized that in seminomas SOX17 uses at least some of the SOX2 binding sites that were described in ESCs to activate pluripotency.

Here we used high-throughput chromatin immunoprecipitation sequencing (ChIP-seq) analysis to investigate SOX17 and SOX2 genome occupancy in seminoma and EC model cell lines to decipher the functional role of both factors in TGCT biology. Based on our results, we present a model where SOX17 is essential for the seminoma cell fate by activating pluripotency-related genes and suppressing differentiation via transactivation of *TFAP2C*, *PRDM1* and *PRDM14*.

## MATERIALS AND METHODS

### Cell Culture

TGCT cell lines were cultured as described previously ^11^.
In brief, TCam-2 (RRID:CVCL_T012) cells (a kind gift from Prof. Dr. J. Shipley, Institute of Cancer Research, Sutton, England) were cultured in RPMI medium (Thermo Fisher Scientific, Waltham, USA). 2102EP (RRID:CVCL_C522) cells (a kind gift from Prof. Dr. L. Looijenga, Erasmus MC, Daniel den Hoed Cancer Center, Rotterdam, Netherlands) were cultured in DMEM medium (Thermo Fisher Scientific). Both media were supplemented with 10% FBS (Thermo Fisher Scientific), 1% penicillin / streptomycin (PAN, Aidenbach, Germany) and 200 mM L-glutamine (PAN). MPAF were obtained from Dr. M. Peitz (Bonn University, Institute of Reconstructive Neurobiology, Bonn, Germany). FS1 Sertoli cells were obtained from Prof. Dr. V. Schumacher (Nephrology Research Center, Boston, USA)^12^. All experiments were performed with mycoplasma-free cells and all cell lines were authenticated using STR analysis.

### Tumour Xenotransplantation

For xenotransplantation, 1 × 10^7^ cells (TCam-2 or 2102EP) were re-suspended in 500 μl of 4°C cold matrigel (Corning, Amsterdam, Netherlands) and injected subcutaneously into the flank of Crl:NU-*Foxn1^nu^* mice (Charles River Laboratories, Erkrath, Germany) as described ^13^. Tumours were isolated six weeks after injection.

### Fixation and Chromatin Preparation

For chromatin preparation from cell lines, 1 × 10^7^ cells / immunoprecipitation were fixed for 30 minutes (min) at room temperature (RT) using Diagenode Crosslink Gold (Diagenode, Liège, Belgium). After two washing steps with PBS (Thermo Fisher Scientific), cells were additionally fixed for 10 min in 1% formaldehyde (Applichem, Darmstadt, Germany) and then processed according to the SimpleChIP® Plus Enzymatic Chromatin IP Kit (Cell Signaling Technology, Leiden, Netherlands). For chromatin preparation from tissue, 25 mg tumour tissue / IP were fixed in 1% formaldehyde (Applichem) and processed according to the SimpleChIP® Plus Enzymatic Chromatin IP Kit (Cell Signaling Technology).

### Chromatin Immunoprecipitation

Chromatin immunoprecipitation (ChIP) was carried out using 200 µg chromatin lysate and 5 µg antibody. The protocol was performed according to the Simple ChIP Enzymatic Chromatin IP Kit (Cell Signaling Technology). 2% Input (= 2 µg chromatin) and IgG-IP served as controls. For ChIP-qPCR 10 µl of IP samples were amplified using the Genomeplex Single Cell Whole Genome Amplification Kit (WGA4) (Sigma Aldrich, Taufkirchen, Germany) and subjected to qPCR in a 1:40 dilution. For primer information, see **Supp. Table 1**.

### Next-Generation Sequencing

ChIP-seq libraries were generated using the MicroPlex Library Preparation Kit v2 (Diagenode). Quality of ChIP-seq libraries was assessed on a 2400 Tapestation (Agilent, Santa Clara, USA) and library pools were quantified using Qubit (Thermo Fisher Scientific). Samples were sequenced on the Illumina High Seq 2500 v4 (Illumina, San Diego, USA) using 25 M single-end reads (1 × 75 bp) per sample. Quality of reads was assessed using FastQC v0.11.8 (https://www.bioinformatics.babraham.ac.uk/projects/download.html#fastqc). ChIP-seq quality statistics are shown in **Supp. Table 2**. Reads were mapped to the human genome (hg38) using CLC Genomics Workbench (Qiagen, Hilden, Germany) and data was analysed using HOMER (Hypergeometric Optimization of Motif EnRichment) Software (http://homer.ucsd.edu/homer/) ^14^. For peak calling, regions were considered significantly enriched where signals in the SOX17 / SOX2 IP were at least four times higher compared to control. Raw data and peak files can be downloaded from gene expression omnibus (GEO): GSE130519. Binding occupancy profiles were calculated using Easeq https://easeq.net/ ^15^. Gene set enrichment analysis (GSEA) on curated gene sets (C2) was performed using the Molecular Signatures Database (MSigDB) http://software.broadinstitute.org/gsea/msigdb/ ^16,17^. Venn diagrams were visualized using Venny2.1 http://bioinfogp.cnb.csic.es/tools/venny/ ^18^. Visualization of transcription factor binding occupancy was performed using the Integrative Genomics Viewer (IGV) browser https://software.broadinstitute.org/software/igv/ ^19,20^. For meta-analysis with previously published SOX17 (GSM1505761, GSM1505762, GSM1505763) and SOX2 ChIP-seq datasets (GSM1505766, GSM1505768), bed-files were downloaded from GEO ^21^.

### CRISPR/Cas9 mediated Deletion of *SOX17* in TCam-2 Cells

To generate *SOX17*-deficient TCam-2 cells (TCam-2^ΔSOX17^), 1×10^5^ cells were transfected with 500 ng pX330-U6-Chimeric_BB-CBh-hSpCas9 (addgene: #42230) coding for *SOX17*-KO-gRNA1 and *SOX17*-KO-gRNA2 (gRNA1: ACGGGTAGCCGTCGAGCGG, gRNA2: GGCACCTACAGCTACGCGC) in a 1:1 ratio. For transfection efficacy monitoring, control cells were transfected with 500 ng pEGFP-N3 (Clontech, # 6080-1). Transfection was performed using FuGENE HD transfection reagent (500 ng DNA: 2.5 µl FuGENE reagent) (Promega, Mannheim, Germany).

### RNA and Protein Isolation

Proteins were isolated using RIPA buffer and protein concentrations were determined using the BCA Protein Assay Kit (Thermo Fisher Scientific). Total RNA was extracted using RNeasy Mini Kit (Qiagen, Hilden, Germany). RNA quality was assessed by photometric measurement of ratios 260/280 nm and 260/230 nm using a NanoDrop photometer (PeqLab, Erlangen, Germany).

### Quantitative real-time RT-PCR

cDNA was synthesized using Maxima First Strand cDNA synthesis Kit (Thermo Fisher Scientific). For qRT-PCR, 7.58 ng of cDNA was run in technical triplicates with Maxima SYBR Green qPCR Master Mix (Fermentas, St. Leon-Rot, Germany). Primer sequences are listed in **Supp. Table 3**. qRT-PCR was performed using the ViiA 7 RealTime PCR System (Thermo Fisher Scientific). *GAPDH* was used as housekeeping gene and for data normalization. Relative expression was calculated from Ct (relative expression = 2^−ΔΔCT^).

### Western blot Analysis

Proteins were separated by SDS gel electrophoresis and transferred onto a PVDF membrane (Carl Roth, Karlsruhe, Germany) using the semi-dry Trans Blot Turbo (Bio-Rad, Düsseldorf, Germany). Membranes were blocked in 5% non-fat milk powder (Applichem) in PBS (Applichem) + 0.05% Tween-20 (Applichem) (PBST). The membrane was then incubated with primary antibody overnight at 4°C, see **Supp. Table 4**. After three washing steps in PBST, the membrane was incubated with HRP-conjugated secondary antibody for 1 hour (h) at RT. After three washing steps in PBST, signal was detected using SuperSignal West Pico Chemiluminescent Substrate (Thermo Fisher Scientific).

### Co-Immunoprecipitation

1.5 mg of Dynabeads Protein G (Thermo Fisher Scientific) were coated with 10 µg of primary antibody (see **Supp. Table 4**), according to the manufacturer’s instructions. After washing the beads-antibody complex with PBS pH 7.4 (+0.02% Tween-20), 1 mg of whole protein lysate were added and incubated with the beads under constant rotation for 30 min at RT. After three washing steps, the beads-antibody-antigen complexes were eluted in 15 µl low pH glycine buffer + 5 µl Roti-Load (Carl Roth) for 5 min at 95°C. The beads were then separated from the antibody-antigen complex in a magnetic rack and the clear supernatant was loaded on a 12% SDS gel for visualization. 20 µg protein lysate served as 2% input sample.

### Immunofluorescence

Cells were washed with PBS (Thermo Fisher Scientific) and fixed in 4% formaldehyde for 10 min at RT. Afterwards, cells were permeabilized using 0.5% Triton X-100 (Applichem) in PBS for 5 min at RT. Cells were washed twice in PBS and blocked in 2% bovine serum albumin in PBS for 1 h at RT. Cells were then incubated with primary antibody (see **Supp. Table 4**) diluted in blocking solution for 2 h at RT or 4°C overnight. After three washing steps in PBS, cells were incubated with Alexa Fluor secondary antibody (Thermo Fisher Scientific) diluted in blocking solution for 1 h at RT or 4°C overnight. Cells were then washed three times in PBS and counterstained with DAPI (Applichem) in PBS for 5 min at RT.

### Alkaline Phosphatase Staining

For detection of alkaline phosphatase (AP) activity, cells were seeded in 6-well dishes. At the desired time point of analysis, cells were fixed for 1 min in 4% formaldehyde and stained for AP activity using the Alkaline Phosphatase Detection Kit (Merck Millipore, Darmstadt, Germany).

### Statistical Analyses

For statistical analyses of qRT-PCR expression data, significance was calculated using two-tailed students t-test (* p ≤ 0.05; ** p ≤ 0.01; *** p ≤ 0.001). qRT-PCR measurements were performed in three biological replicates, where error bars indicate standard deviation from the mean. For statistical analyses of motif enrichment within ChIP-seq data, p-values were calculated using Homer Software (http://homer.ucsd.edu/homer/) ^14^.

## RESULTS

### Binding Sites of SOX17 and SOX2 in TGCT Cells

We hypothesized that in seminoma cells SOX17 and OCT4 partner to regulate pluripotency genes, while SOX17-binding to endodermal genes might be inhibited. By co-immunoprecipitation we verified interaction of SOX17 and OCT4 in seminoma-like TCam-2 cells (**Supp. Fig. 1A-B**), which had already been demonstrated in our previous study ^22^. In ESCs, SOX2/OCT4 binds to canonical motifs to regulate pluripotency genes. Because EC cells are reminiscent of ESCs in expressing pluripotency genes we predicted a similar SOX2-OCT4 interaction in EC cells. Using co-immunoprecipitation we demonstrated interaction of SOX2 and OCT4 in the EC line 2102EP (**Supp. Fig. 1C-D**). Next, suitability of SOX17 and SOX2 antibodies for ChIP was tested by immunoblotting of SOX17- and SOX2-precipitated chromatin lysates. We detected a band indicative of the SOX17 protein in the SOX17 ChIP sample and SOX2 protein in the SOX2 ChIP sample (**Supp. Fig. 1E**), confirming that SOX17 and SOX2 were successfully immunoprecipitated.

To determine genome occupancy of SOX17 and SOX2 in TGCT cell lines, we performed ChIP-seq analysis in TCam-2 (SOX17) and 2102EP (SOX2) cells on three independent biological replicates. We detected a total of 17,509 SOX17 peaks (present in all three replicates) in TCam-2 (**Supp. Data 1A-B upon request, Fig. 1A**) and 17,797 SOX2 peaks (present in all three replicates) in 2102EP (**Supp. Data 1C-D upon request, Fig. 1A**). Interestingly, 4,427 of SOX17 and SOX2 peaks overlapped, indicating common binding sites of SOX17 and SOX2 in TCam-2 and 2102EP, respectively (Fig. 1A). Comparison of SOX17 and SOX2 genome binding occupancy in TGCT cells (+/− 10,000 bp from transcription start site (TSS)), confirms that ChIP-seq signals correlate to each other (r = 0.633) (Fig. 1B). SOX17 and SOX2 binding occupancy +/− 10,000 bp from TSS further illustrates highest density in a region spanning +/− 1,500 bp from the TSS, an effect that was even more prominent for SOX17 binding in TCam-2 cells (Fig. 1C; dashed line). This indicates that both transcription factors mainly bind in close proximity to promoter regions at a similar distance to the TSS.

**Figure 1:**
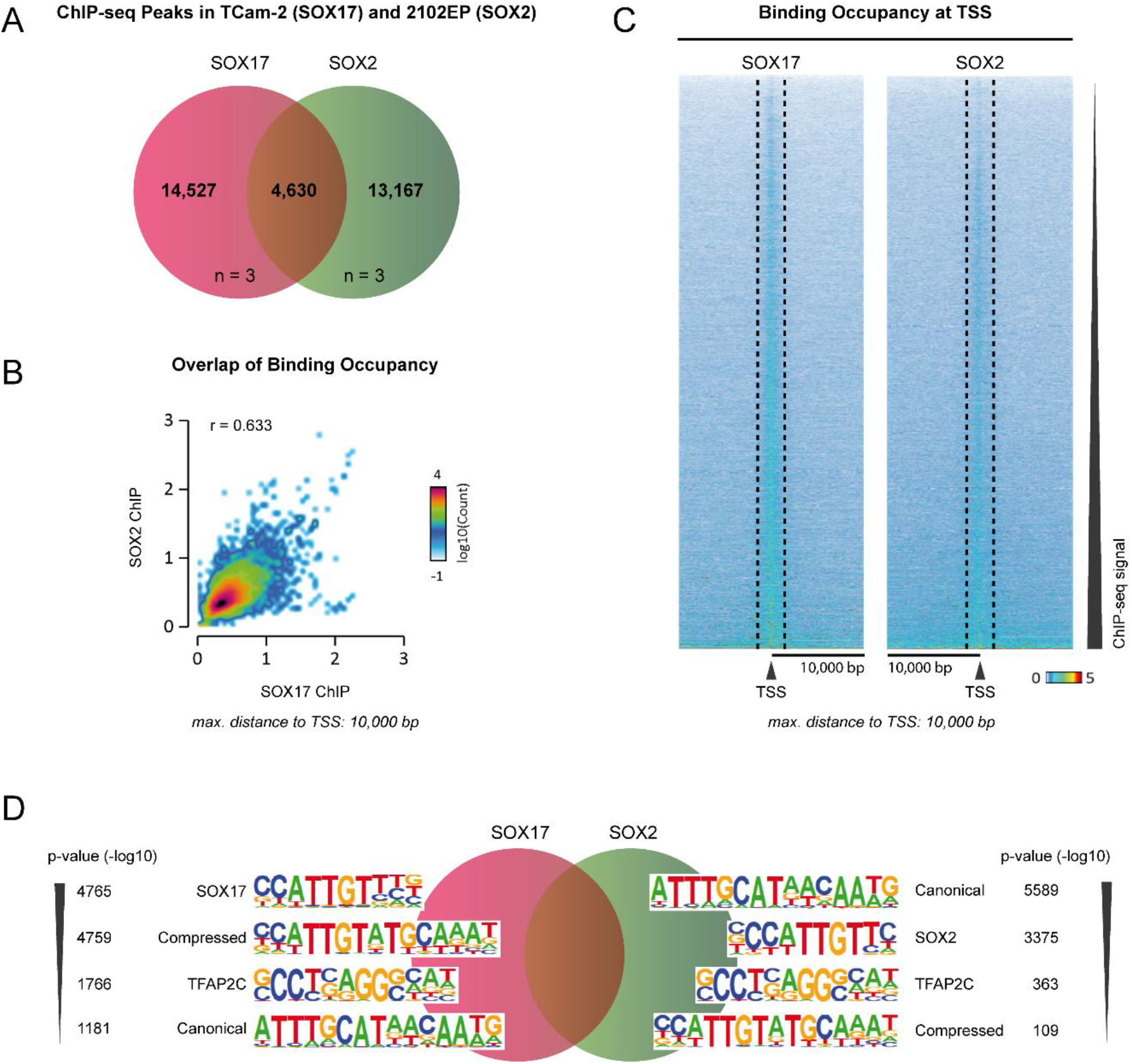
**(A)** Venn diagram depicting the number of annotated peaks identified commonly in all three biological replicates (n = 3) of SOX17 and SOX2 ChIP-seq samples. Red circles = SOX17 ChIP-seq analyses in TCam-2 cells. Green circles = SOX2 ChIP-seq analyses in 2102EP cells. **(B)** 2D histogram of the SOX17 ChIP (on x-axis) and SOX2 ChIP (on y-axis) signals found +/− 10,000 bp from TSS of genes with correlation coefficient (r). **(C)** Heat map of SOX17 and SOX2 ChIP signal 10,000 bp upstream and downstream of the transcription start site (TSS) of genes. Regions were sorted according to decreasing SOX17 or SOX2 ChIP signal. Dashed lines are spanning regions 1,500 bp +/−TSS. **(D)** Transcription factor binding motifs enriched in SOX17 and SOX2 ChIP-seq datasets, determined by HOMER ^13^. The relative enrichment to background sequences was calculated as indicated (p-value).

We next performed HOMER analysis to identify motifs in promoter and genomic regions directly bound by SOX17 in TCam-2 and SOX2 in 2102EP cells. As expected, in TCam-2 cells regions bound by SOX17 showed strongest enrichment for the SOX17 binding motif (-log10 (p-value): 4,606) and the compressed motif (-log10 (p-value): 4,533) (Fig. 1D, **Supp. Data 2A upon request**). Additionally, SOX17 recognizes and binds related SOX-family binding motifs (SOX3, SOX15, SOX10, SOX2, SOX6, SOX4 and SOX9) (**Supp. Data 2A upon request**). Further, within SOX17-bound regions, we detected TFAP2C (AP2-gamma) binding motifs (-log10 (p-value): 1,555) (Fig. 1D, **Supp. Data 2A upon request**). TFAP2C is highly expressed in seminomas ^3^. This suggests that SOX17 and TFAP2C might cooperate to regulate a set of common target genes in TCam-2. Interestingly, SOX17 peaks also mapped to canonical motifs (-log10 (p-value): 1,072) (Fig. 1D, **Supp. Data 2A upon request**). In 2102EP, regions bound by SOX2 showed strongest enrichment for the canonical motif (-log10 (p-value): 5,589) (Fig. 1D, **Supp. Data 2B upon request**). Further, we found enrichment of SOX3, SOX2, SOX6, SOX10, SOX15, SOX17, SOX4 and SOX9 binding motifs (**Supp. Data 2B upon request**). Also in 2102EP cells, which express TFAP2C at moderate levels, SOX2-bound regions were enriched for the TFAP2C motif (-log10 (p-value): 363) (Fig. 1D, **Supp. Data 2B upon request**). Interestingly, only a minor fraction of SOX2 peaks displayed the compressed motif (-log10 (p-value): 109) (Fig. 1D).

Additionally, we analysed binding motifs identified in those regions that were commonly bound by SOX17 in TCam-2 and SOX2 in 2102EP cells. Expectedly, common DNA binding sites were similarly enriched for SOX motifs and the TFAP2C motif (**Supp. Data 2C upon request**). Strong enrichment was also detected for the canonical motif (-log10 (p-value): 1,083) (**Supp. Data 2C upon request**). Lower enrichment was seen for the compressed motif (-log10 (p-value): 191) (**Supp. Data 2C upon request**). Taken together, these findings illustrate that SOX17 peaks in TCam-2 and SOX2 peaks in 2102EP mostly overlap at canonical and SOX-family binding motifs.

### The Genes regulated by SOX17 and SOX2 in TGCT cells

We next annotated the SOX17 and SOX2 peaks to genes with the most closely located TSS (only in a region +/− 10,000 bp from TSS). In TCam-2 cells, this included 3,461 genes bound by SOX17 and in 2102EP cells 3,269 genes bound by SOX2. Gene set enrichment analysis of both datasets revealed a significant overlap with SOX2 and NANOG target genes (**Supp. Fig. 2A-B**), confirming their role in regulating pluripotency-associated genes. Since in TGCTs both SOX17 and SOX2 share target genes with NANOG, we analysed whether there is a direct protein-protein interaction. However, co-immunoprecipitation showed no interaction of SOX17 or SOX2 with NANOG (**Supp. Fig. 2C**).

We then meta-analysed SOX17- and SOX2-bound genes with previously published gene expression data of TCam-2 and 2102EP, respectively ^23–25^. In TCam-2 cells, we identified 31 genes above an arbitrary expression threshold of ≥ log_2_7 and SOX17 peak score of ≥ 150 (Fig. 2A). We discovered that SOX17 is a regulator of the insulin-like growth factor *IGF1*, which is known to be overexpressed in seminoma and TCam-2 cells ^26^. Further, we found that 14 of these 31 genes (*NANOG, ZNF511, GLDC, PLEKHG6, XPNPEP1, C8orf76, TFDP2, LRRC59, MIR205, PSAP, LOC728715, SRGAP1, KNTC1, AACS*) were exclusively bound by SOX17 in seminoma cells, but not in endoderm-, mesoderm- or mesendoderm-differentiated ESCs ^27^ (**Supp. Data 3A upon request**). This suggests that these genes contribute to the germ cell fate and / or latent pluripotency of seminomas. Of these, the pluripotency factor NANOG is a well-established marker of seminomas and ECs ^28^. Not surprisingly, we therefore found *NANOG* also among SOX2 target genes in 2102EP cells (**Supp. Data 1D upon request**).

**Figure 2:**
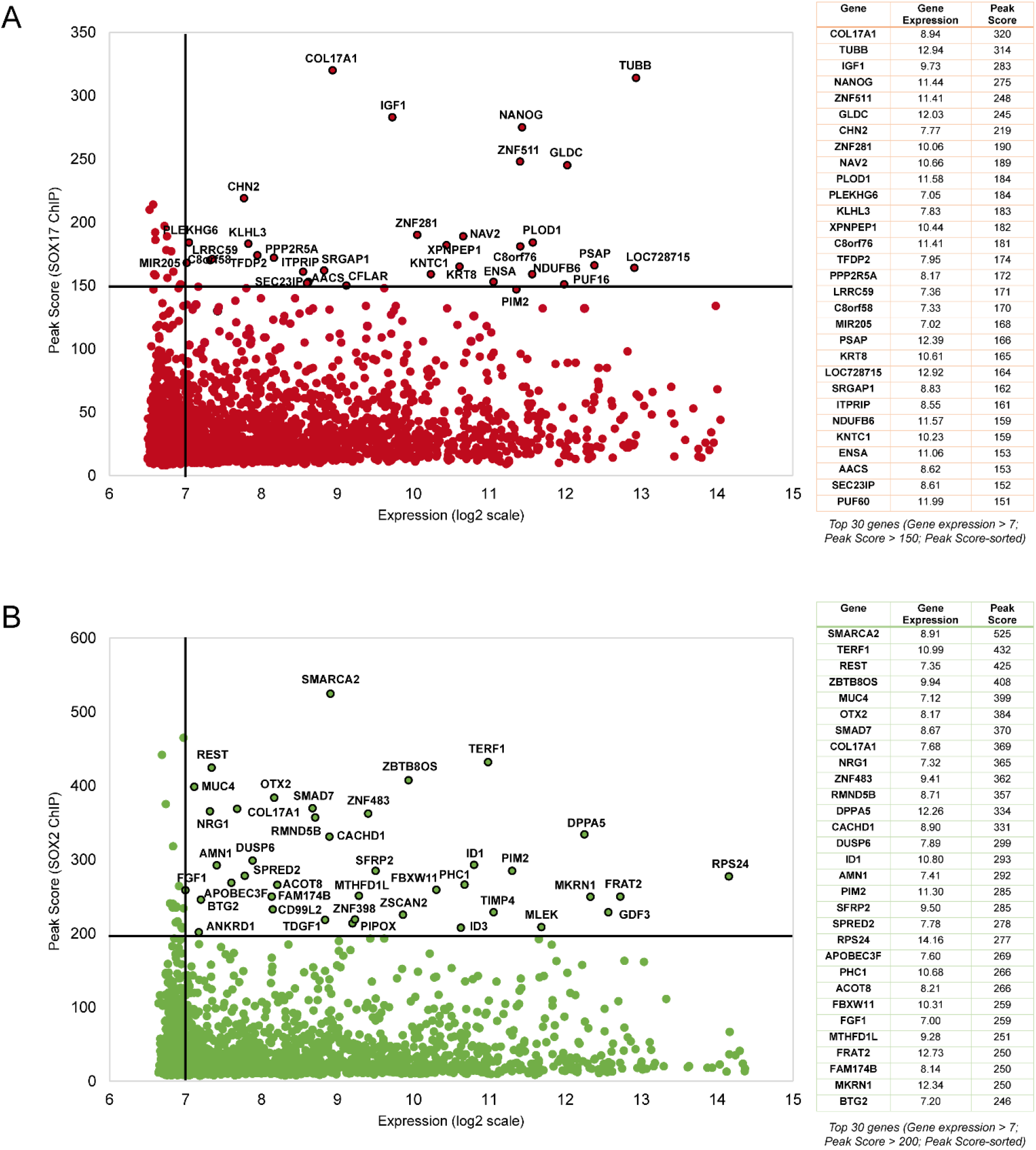
**(A)** Left: Scatterplot of genes, which are bound by SOX17 within a distance +/− 10,000 bp from TSS. The relative gene expression of each gene in TCam-2 cells is given on the x-axis, SOX17 ChIP-seq peak score is given on the y-axis. Right: Table of top 30 genes (peak-score-sorted) with a gene expression > 7. **(B)** Left: Scatterplot of genes, which are bound by SOX2 within a distance +/− 10,000 bp from TSS. The relative gene expression of each gene in 2102EP cells is given on the x-axis, SOX2 ChIP-seq peak score is given on the y-axis. Right: Table of top 30 genes (peak-score-sorted) with a gene expression > 7.

In 2102EP cells we identified 40 genes above an arbitrary gene expression threshold of ≥ log_2_7 and SOX2 peak score of ≥ 200 (Fig. 2B). Of these*, GDF3, TDGF1* and *DPPA5* are well-known regulators of pluripotency ^29–31^. Further, *OTX2* was described as a gatekeeper of PGC specification in mice ^32^. Previously published microarray data reveals a higher expression of *OTX2* in TGCT tissues compared to normal testis tissue ^33^ (**Supp. Fig. 2D**). *OTX2* was also identified as a direct target of SOX17 in TCam-2 cells (**Supp. Data 1B upon request**). We then compared EC to ESC in order to find genes, which are linked to tumorigenesis. Comparing our SOX2 ChIP-seq data from 2102EP to data from ESC (HUE64) ^27^ revealed that four of the 40 genes (*MLEC, PIM2, CD99L2*, APOBEC3F) were exclusively bound by SOX2 in 2102EP (**Supp. Data 3B upon request**). *PIM2* is a proto-oncogene regulating survival and proliferation ^34^. We found that *PIM2* is highly expressed in 2102EP and TCam-2 cells, but weakly in the Sertoli cell line FS1 and adult fibroblasts (MPAF) (**Supp. Fig. 2E**). This confirms findings from Jiménez-García *et al*., who demonstrated overexpression of *PIM1* and *PIM2* in TGCTs ^34^. Interestingly, *PIM2* was also bound by SOX17 (peak score: 146) in TCam-2 cells (Fig. 2A), suggesting that SOX2 and SOX17 are able to transactivate *PIM2*. Notably, *PIM1* was similarly bound by SOX17 and SOX2 in TGCT cells, albeit with a lower enrichment (SOX17 peak score: 31; SOX2 peak score: 21) (**Supp. Data 1B,D upon request**). Together, these data demonstrate that known markers of seminomas (*IGF1, NANOG, PIM1/2*) and / or ECs (*GDF3, TDGF1, NANOG, PIM1/2*) are under direct control of SOX17 and SOX2, respectively. Additionally, we present *OTX2*, which is overexpressed in TGCTs, as a *bona fide* target of SOX17 and SOX2.

### SOX17 and SOX2 regulate Germ Cell- and Pluripotency-associated Genes in TGCTs

Next, we asked whether we could find additional genes under control of SOX17 in TCam-2 and SOX2 in 2102EP that are related to germ cell (GC) development, cancer and / or pluripotency. We found that SOX17 in TCam-2 binds and possibly regulates the cancer-related gene *MYC* ^35^, the GC specifier *PRDM1* ^36^ and the BMP member *BMP7*, known to contribute to GC differentiation in mice ^37^ (Fig. 3A). SOX2-bound genes in EC cells included regulators of proliferation, cancer and pluripotency (*GDF3, LEFTY2, SALL4, SOX2, TP53, POU5F1 (OCT4)*) (Fig. 3A). Comparison of SOX17 and SOX2-bound genes in TCam-2 and 2102EP indicates that 1,268 genes overlap, representing approximately one third of all regulated genes of each factor (Fig. 3A). Among these, we found key-regulators of GC development and / or pluripotency, such as *PRDM14*, *DPPA4*, *TDGF1*, *LIN28A*, and *TRIM71* (in addition to *PIM1/2*, *OTX2* and *NANOG)* (Fig. 3A). All of these genes are highly expressed in TCam-2 and 2102EP, except for *TDGF1*, which is expressed in 2102EP only (**Supp. Fig. 3A**). Next, we asked which DNA motifs SOX17 and SOX2 use to regulate these genes. Canonical motifs were found within regulatory regions of *PIM2*, *TRIM71*, *PRDM14, GDF3* and *SOX2*, while compressed motifs were found in *PRDM1, MYC* and *IGF1* (Fig. 3B-E). The other SOX17 and SOX2 target genes (*PIM1, OTX2, DPPA4, TDGF1, NANOG* and *LIN28A*) seem to be regulated independent of OCT4, since neither compressed nor canonical motifs were detected (Fig. 3B - E).

**Figure 3:**
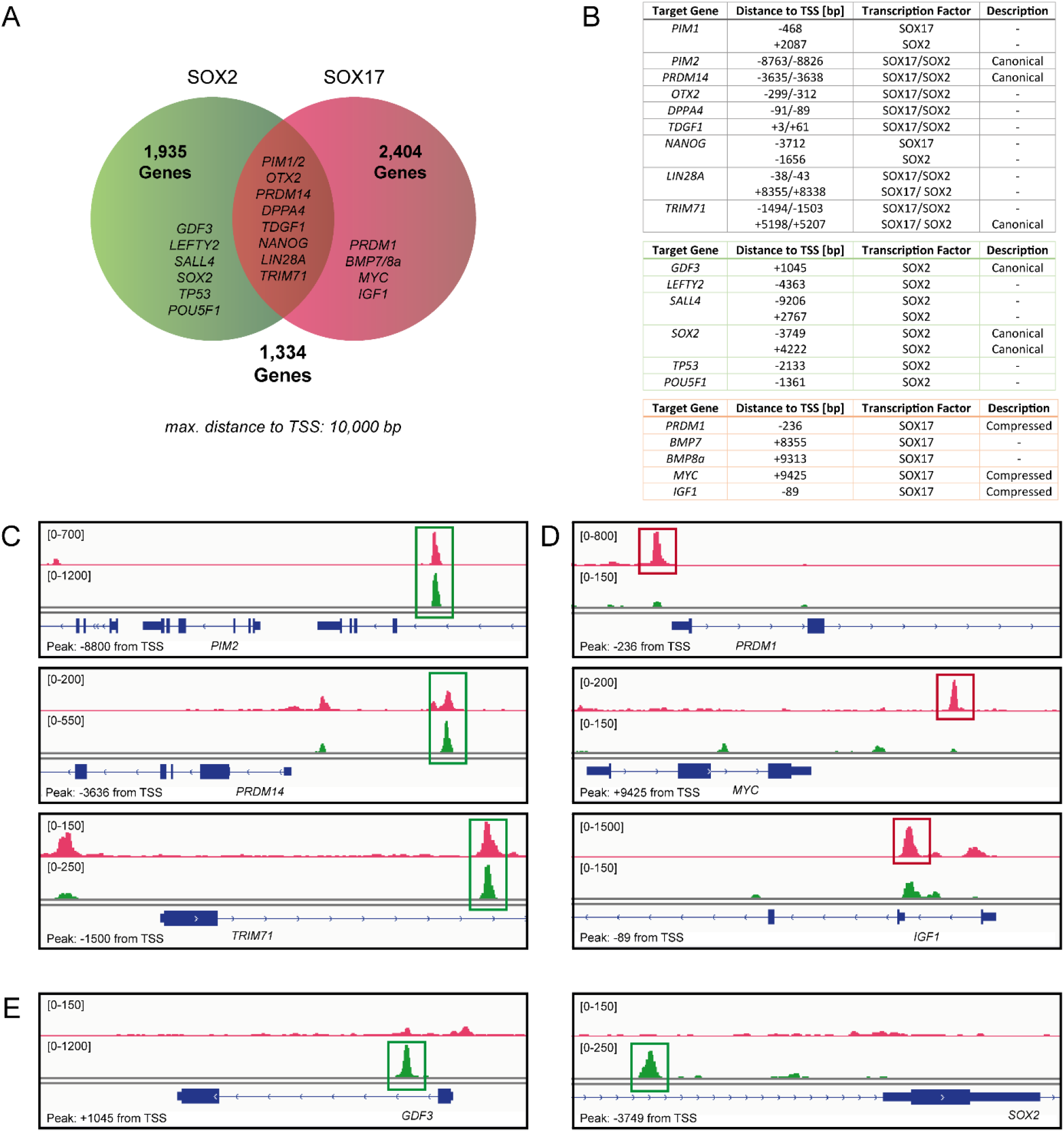
**(A)** Venn diagram depicting the number of genes bound by SOX17, SOX2 or SOX17 and SOX2 (binding site within a distance +/− 10,000 from TSS) in TCam-2 and 2102EP, respectively. Genes representative for each group are listed. **(B)** Description of binding sites (relative to TSS) of SOX17 and SOX2 in the regulatory regions of the genes listed in (A). **(C-E)** DNA binding profiles of SOX17 and SOX2 at canonical motifs (green boxes) of *PIM2*, *PRDM14*, *TRIM71* **(C)**, at compressed motifs (red boxes) of *PRDM1*, *MYC*, *IGF1* **(D)** and at canonical motifs (green boxes) of *GDF3*, *SOX2* **(E)**. Distance of binding sites to TSS are indicated in lower left corner. Scales (upper left corner) were adjusted to allow for best comparison.

In order to confirm that the SOX2 regulatory network is also established and maintained *in vivo*, we xenografted 2102EP and TCam-2 cells into the flank of nude mice and isolated tumours after six weeks. 2102EP xenografts formed compact EC tumours, while TCam-2 cells reprogrammed to an EC-like state, as published previously ^23^. Both tumours highly expressed SOX2 and were subjected to SOX2 ChIP-qPCR. Similar to 2102EP cells *in vitro*, we were able to detect strong enrichment of SOX2 in the regulatory regions of *DPPA4*, *LEFTY2*, *LIN28A*, *TDGF1, NANOG, GDF3, TP53, PRDM14, SALL4* and *SOX2* in 2102EP tumours (n = 4) (**Supp. Fig. 4A**). Although less pronounced, TCam-2-derived reprogrammed EC-like tumours similarly demonstrate enrichment of SOX2 in the described regulatory regions in at least two of three samples (**Supp. Fig. 4B**).

### Germ cell-related Function of SOX17 in Seminoma Cells

Next, we asked how the role of SOX17 in seminomas is different to the role of SOX17 as endodermal factor. Detailed analysis revealed that from 3,461 genes bound by SOX17 in TCam-2 (10,000 bp +/− from TSS), 66% are exclusively found in TCam-2, but not in mesoderm-, endoderm- or mesendoderm-differentiated ESCs (10,000 bp +/− from TSS) ^27^ (Fig. 4A). Even more strikingly, 89% of the *bona fide* SOX17 binding sites identified in TCam-2 were not bound by SOX17 in differentiated ESCs ^27^ (Fig. 4A). This shows that the DNA binding sites of SOX17 differ dramatically between TCam-2 and somatic lineages. The most prominent difference is that in TCam-2 we found strong enrichment of SOX17 at the canonical motif, which was absent in somatic cells (Fig. 4B). Surprisingly, also enrichment for the compressed motif was much stronger in TCam-2 cells compared to differentiated ESCs (Fig. 4B). This shows that SOX17 fulfills its function in seminoma cells not only via the canonical, but also the compressed motif.

**Figure 4:**
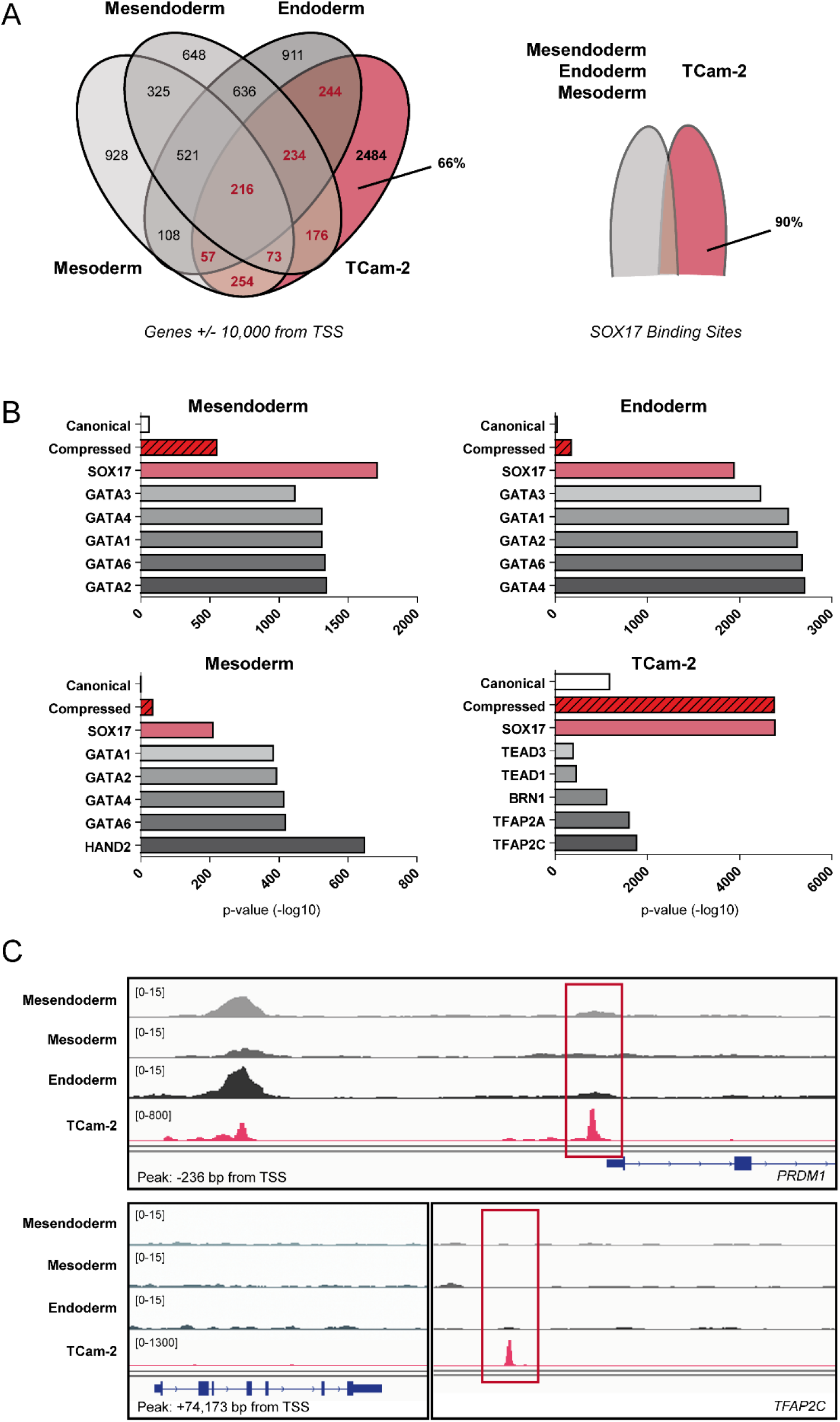
**(A)** Left: Venn diagram depicting the number of genes putatively regulated by SOX17 (SOX17 binding site within a distance +/− 10,000 from TSS) in mesendoderm-, endoderm- and mesoderm-differentiated ESCs, as well as TCam-2. Right: Venn diagram depicting the overlap of SOX17 binding sites in TCam-2 cells with SOX17 in mesendoderm-, endoderm- and/or mesoderm-differentiated ESCs. **(B)** Transcription factor binding motifs enriched within SOX17 ChIP-seq peaks of mesendoderm-, endoderm- and mesoderm-differentiated ESCs, as well as TCam-2, as determined by HOMER ^13^. **(C)** Upper panel: DNA binding profile of SOX17 upstream of *PRDM1* in mesendoderm-, endoderm- and mesoderm-differentiated ESCs, as well as TCam-2. Red box indicates the compressed motif bound by SOX17 in TCam-2 cells. Lower panel: DNA binding profile of SOX17 downstream of *TFAP2C* in mesendoderm-, endoderm- and mesoderm-differentiated ESCs, as well as TCam-2. Red box indicates the compressed motif bound by SOX17 in TCam-2 cells. Distance of binding sites to TSS are indicated in lower left corner. Scales (upper left corner) were adjusted to allow for best comparison.

But how does the role of SOX17 in seminomas compare to the role of SOX17 in PGCs? Previous studies demonstrated that human PGC specification is governed by a transcription factor network of SOX17, TFAP2C and PRDM1, where SOX17 transactivates *TFAP2C* and *PRDM1* ^10,38,39^. In TCam-2, we found that SOX17 binds to a compressed motif within the *PRDM1* promoter, which is not bound by SOX17 in somatic lineages (Fig. 4C). Surprisingly, in TCam-2 *TFAP2C* was not bound by SOX17 +/− 10,000 bp from TSS. However, we found in total five binding sites of SOX17 in distal regulatory regions (−19,773; +74,173; +124,640; +175,507; +268,206) upstream and downstream of the *TFAP2C* TSS (**Supp. Data 1B upon request**). The highest ChIP-seq peak score was calculated for a compressed motif, which lies closest to the *TFAP2C* TSS (+74,173). Again, this binding site was not bound by SOX17 in somatic lineages (Fig. 4C). Thus, in seminomas the transcription factor network (SOX17-PRDM1-TFAP2C) described in human PGC specification seems to be in place, but absent in differentiated ESCs.

### Loss of SOX17 in Seminoma Cells results in Differentiation

To test whether SOX17 regulates latent pluripotency of seminomas, we used CRISPR/Cas9-mediated gene-editing to delete a region of approximately 130 bp within the *SOX17* coding sequence (Fig. 5A). *SOX17* is encoded on chromosome ^8^, which has six copies in TCam-2 cells ^40^. Therefore, gene editing of the *SOX17* locus in TCam-2 cells results in a mixed population of cells having one to six alleles of *SOX17* deleted. The loss of SOX17 protein in the gene-edited TCam-2 bulk population (referred to as TCam-2^ΔSOX17^) was shown by Western blot analysis (Fig. 5B). TCam-2^ΔSOX17^ cells show morphologically distinct areas suggestive of differentiation (Fig. 5C). They display polynucleated trophoblast giant cell morphology (Fig. 5D), which is reminiscent of TCam-2 cells differentiated by treatment with TGFβ, EGF and FGF4 ^41^. In these morphologically distinct areas, TCam-2^ΔSOX17^ are negative for alkaline phosphatase (AP) activity, indicating loss of pluripotency (Fig. 5E). qRT-PCR analysis confirmed a downregulation of pluripotency and / or seminoma markers (*POU5F1, TFAP2C, NANOG* and *LIN28B* (but not *LIN28A*)) in TCam-2^ΔSOX17^ cells (Fig. 5F). Immunofluorescence staining confirmed reduction of OCT4 and TFAP2C protein levels in those cells that display reduced levels or complete loss of SOX17 protein (Fig. 5G). Further, we detected downregulation of additional GC- and / or pluripotency-associated markers (*PRDM14, PRDM1, KIT, NANOS3, SPRY4, ALPL*), independent of whether they were identified as direct SOX17 targets or not (**Supp. Fig. 5A**).

**Figure 5:**
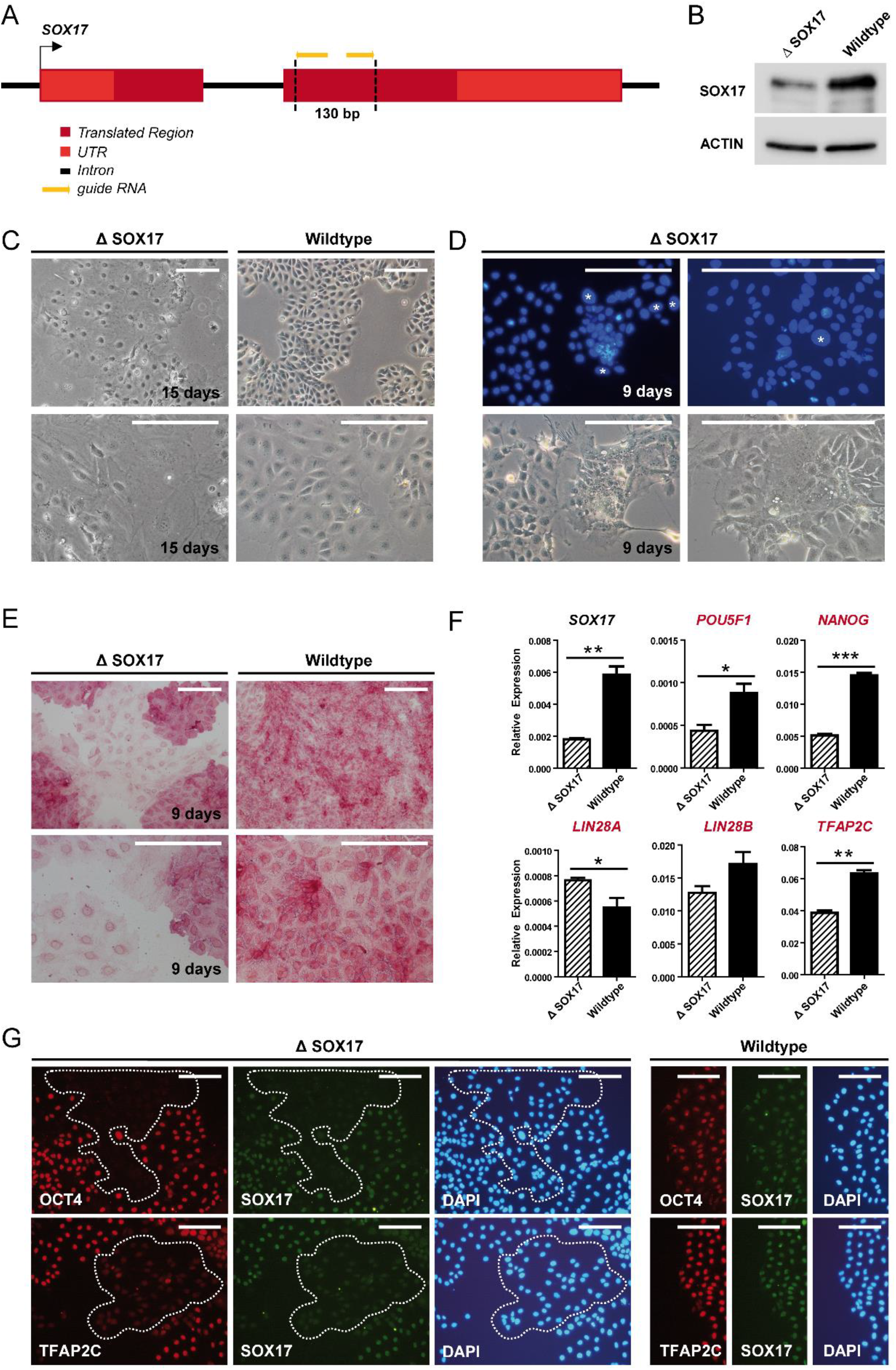
**(A)** CRISPR/Cas9-gene editing strategy of SOX17 locus in TCam-2 cells. **(B)** Immunoblot of SOX17 in gene-edited TCam-2^ΔSOX17^ cells. β-ACTIN (ACTIN) served as loading control. **(C)** Morphology of TCam-2^ΔSOX17^ cells (left) and TCam-2 wildtype cells (right) 15 days following gene-editing. Scale bar = 250 µm. **(D)** Hoechst staining of TCam-2^ΔSOX17^ cells (upper panel) and corresponding brightfield images (lower panel) 9 days following gene-editing. Multi-nucleated cells are indicated with an asterisk. Scale bar = 250 µm. **(E)** Alkaline phosphatase staining of TCam-2 wildtype cells and TCam-2^ΔSOX17^ cells 9 days following gene-editing. Scale bar = 250 µm. **(F)** qRT-PCR of selected germ cell markers and SOX17 target genes (red labels) in TCam-2^ΔSOX17^ cells and TCam-2 cells transiently transfected with a GFP-coding plasmid as control (wildtype), 72 h following transfection. Expression was normalized to *GAPDH* as housekeeper. Significance was calculated using two-tailed students t-test (* p ≤ 0.05; ** p ≤ 0.01; *** p ≤ 0.001). Error bars indicate standard deviation from the mean. **(G)** Immunofluorescence staining of OCT4, TFAP2C and SOX17 protein in TCam-2^ΔSOX17^ cells and TCam-2 wildtype cells. Cell nuclei were stained with DAPI. Scale bar = 125 µm.

Next, we asked into which lineage or cell type TCam-2^ΔSOX17^ differentiate. Since SOX17 is known as a master regulator of the endodermal lineage, differentiation into the endoderm lineage seems highly unlikely. The fact that endodermal-differentiation factors *FOXA1* or *FOXA2* were not upregulated further supported this notion (**Supp. Fig. 5B**) 42. In addition, mesoderm or ectoderm differentiation markers were not upregulated (except for mild induction of *PAX6*), nor was SOX2 (**Supp. Fig. 5B**). Therefore, TCam-2 cells do not differentiate into somatic lineages upon loss of SOX17. Due to the resemblance of TCam-2^ΔSOX17^ cells to trophoblast giant cells we screened for expression of extra-embryonic lineage markers. Loss of SOX17 resulted in significant upregulation of the trophoblast giant cell marker *HAND1*, as well as *GATA3, GATA6* and *αHCG* (**Supp. Fig. 5C**). *CDX2* and *EOMES* levels remained unchanged (**Supp. Fig. 5C**). Upregulation of GATA3 was confirmed by immunofluorescence staining (**Supp. Fig. 5D**). Together this suggests that downregulation of SOX17 results in extra-embryonic differentiation.

## DISCUSSION

Here, we have investigated the binding-patterns and target genes of SOX17 and SOX2 in seminoma and EC cell lines, respectively. We demonstrated that both transcription factors bind to canonical motifs by partnering with OCT4, as well as to SOX-family DNA binding motifs. This way SOX17 and SOX2 regulate a common set of pluripotency and germ cell-related genes (*PRDM14, DPPA4, TDGF1, NANOG, LIN28A, TRIM71, OTX2, PIM2*) (Fig. 6). Additionally, in TCam-2 cells SOX17 binds to compressed motifs or SOX motifs (not bound by SOX2 in ECs), thereby regulating the PGC specifiers *PRDM1* and *TFAP2C*, the GC-related BMP member *BMP7* and the cancer-related genes *MYC* and *IGF1* (Fig. 6). In 2102EP EC cells, SOX2 further binds canonical elements or SOX motifs (not bound by SOX17 in TCam-2), thereby regulating additional pluripotency genes (*GDF3, LEFTY2, SALL4, SOX2* and *POU5F1*) (Fig. 6). In contrast to SOX17, SOX2 did not bind to compressed motifs. This is in line with the previous observation of Jauch *et al.* that due to sterical hindrance the SOX2/OCT4 complex is occluded from the compressed motif ^43^. Our findings suggest that in seminomas and probably PGCs, SOX17 is able to functionally replace SOX2 in its role in maintaining pluripotency.

**Figure 6:**
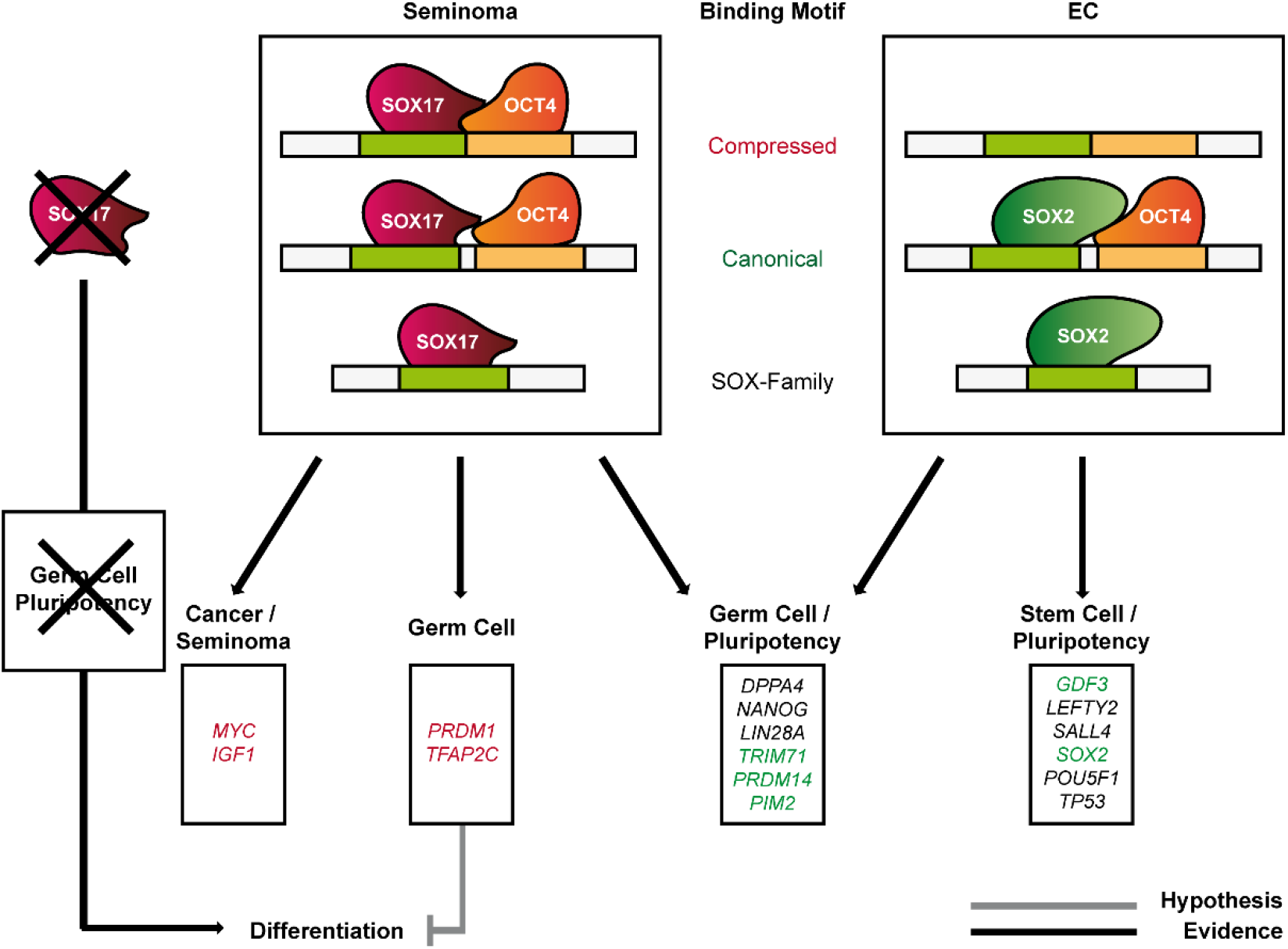
Schematic representation of the role of SOX17 in seminomas (left) and SOX2 in ECs (right). Genes regulated via compressed motifs are written in red labels, genes regulated via canonical motifs are written in green labels. Genes regulated via solitaire SOX motifs are written in black labels.

The role of SOX2 in transcriptional regulation was already well-described for ESCs. Here, we have shown that SOX2 in EC cells regulates similar target genes as SOX2 in ESCs, hence it is a key regulator of pluripotency genes. In addition to already well-established markers of pluripotency, we found *PIM2* and *OTX2* to be SOX2 target genes in ECs. Overexpression of *PIM1* and *PIM2* in mice resulted in an increase in inflammatory signatures as well as an overall higher cell proliferation rate in testicular tissues ^34^. Both *PIM2* and *OTX2* are overexpressed in TGCT, but not much is known about their role in human TGCTs. So, future studies will address the role of *PIM1/2* and *OTX2* for human TGCT development in more detail.

SOX17 is known as a regulator of endodermal cell fate decisions in ESCs and as a key specifier of human PGC fate. Here we demonstrate that only 11% of the SOX17 binding sites in TCam-2 are shared by SOX17 binding sites in differentiated (endoderm-, mesoderm-, mesendoderm-) lineages. The canonical motif was not found among SOX17 binding elements in differentiated lineages. This suggests that SOX17 binding to canonical motifs is specific to seminoma cells (and possibly PGCs). This is a highly important finding, as it demonstrates that the DNA-occupancy of SOX17 is strictly dependent on the cell type analysed. We speculate that either cofactors and / or locus accessibility regulated by epigenetic changes such as DNA methylation or histone modifications might guide SOX17 to its respective target gene. Since seminoma cells highly express pluripotency-associated genes, we predicted that SOX17 transactivates these genes via the canonical motif ^22^. Indeed, we found that in seminoma cells SOX17 binds to canonical motifs allocated within regulatory regions of pluripotency genes (such as *PRDM14, TRIM71*). Additionally, SOX17 binds compressed motifs within the regulatory regions of GC- and cancer-associated genes (such as *IGF1, MYC, TFAP2C, PRDM1, PRDM14*). Therefore, partnering with OCT4 is required in order to fulfil the GC- and cancer-specific function of SOX17 in seminoma cells and to maintain latent pluripotency (Fig. 6).

Thus, the role of SOX17 in seminomas seems to be reminiscent of the role of SOX17 in human PGCs, which is to maintain latent pluripotency and to suppress somatic differentiation via transactivation of *TFAP2C* and *PRDM1*. In human PGCs, SOX17, TFAP2C and PRDM1 form a tripartite transcription factor network, which prohibits mesoderm and neural differentiation and maintains expression of pluripotency- and germ cell-associated genes. Irie *et al.* proposed SOX17 as the most upstream transcription factor within this network ^10^. Later, Kobayashi *et al*. showed that SOX17 and TFAP2C have a pivotal role in specification of human PGCs and that SOX17 stimulates *PRDM1* expression ^36^. In the end, in PGC-like cells all three transcription factors form a self-regulatory network including feedback loops regulating their own expression ^36,39^. Interestingly, PRDM14 was recently introduced as a fourth factor (next to SOX17, TFAP2C and PRDM1) maintaining human PGC pluripotency ^44^. A different study further suggested that an aberrant overexpression of *PRDM14* during PGC development could favour a malignant transformation of the cells ^45^. We here demonstrate that in seminoma cells SOX17 binds multiple elements within *PRDM14* regulatory regions. Thus, SOX17 not only regulates the GC network in seminoma cells by induction of *PRDM1* and *TFAP2C*, but additionally via *PRDM14* expression. Additionally, SOX17 contributes to the latent pluripotency state of seminoma cells by directly transactivating a set of pluripotency-related genes (such as *DPPA4, NANOG, LIN28A, TRIM71*) (Fig. 6). Collectively, these findings highlight the role of SOX17 in the germline and demonstrate the dependency of seminoma cells for SOX17-driven transactivation of pluripotency- and GC-associated genes.

## LIST OF ABBREVIATIONS

TGCT: Testicular Germ Cell Tumour
EC: Embryonal Carcinoma
GCNIS: Germ Cell Neoplasia in Situ
PGC: Primordial Germ Cell
ESC: Embryonic Stem Cell
TSS: Transcription Start Site
GC: Germ Cell
GSEA: Gene Set Enrichment Analysis
GEO: Gene Expression Omnibus
AP: alkaline phosphatase

## ACKNOWLEDGEMENTS

We kindly thank Blanca Randel, Gaby Beine, Angela Egert and Andrea Jäger for technical assistance. Further, we thank the NGS Core Facility of the University Hospital Bonn for the generation of the sequencing data and the Core Unit for Bioinformatic Analysis of the University Hospital Bonn for their assistance with the data analysis.

