## Supplementary Figures and Tables for "Unique and redundant roles of SOX2 and SOX17 in regulating the germ cell tumour fate"

Supp. Figure 1

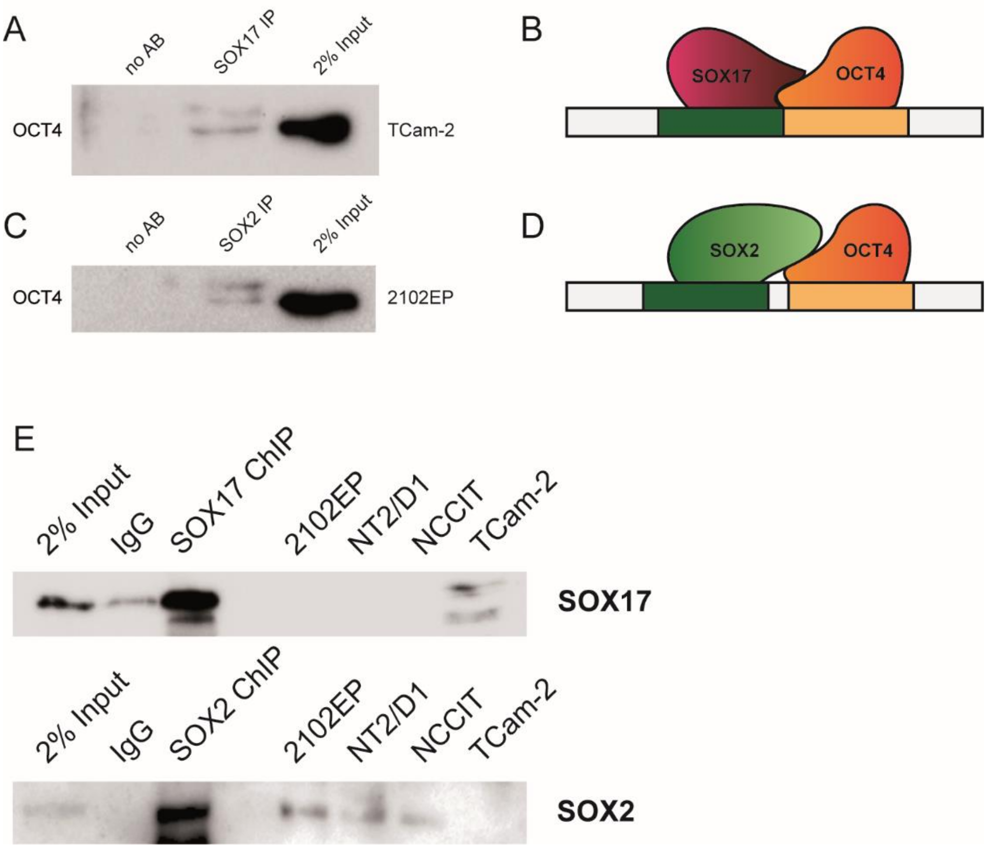


**Supplementary Figure 1:** (A) Co-immunoprecipitation of SOX17 (in TCam-2 cells) with OCT4. 2% input and no-antibody IP served as positive and negative controls, respectively. (B) Graphical illustration of SOX17 partnering with OCT4 to bind to compressed binding motifs. Adapted from 43. (C) Co-immunoprecipitation of SOX2 (in 2102EP cells) with OCT4. 2% input and no-antibody IP served as positive and negative controls, respectively. (D) Graphical illustration of SOX2 partnering with OCT4 to bind to canonical binding motifs. Adapted from 43. (E) Immunoblot of SOX17 and SOX2 protein in SOX17 (TCam-2) and SOX2 (2102EP) ChIP samples. 2% input and IgG IP served as positive and negative controls, respectively. Protein lysates of EC cells (2102EP, NCCIT, NT2/D1) and seminoma cells (TCam-2) served as controls.

### Supp. Figure 2

A

| Gene Set Name [# Genes (K)] | # Genes in Overlap (k) | p-value |
|---|---|---|
| BENPORATH_**SOX2**_TARGETS [734] | 197 | $1.45\ e^{-83}$ |
| BENPORATH_**NANOG**_TARGETS [988] | 228 | $1.12\ e^{-82}$ |
| BLALOCK_ALZHEIMERS_DISEASE_UP [1691] | 290 | $2.79\ e^{-73}$ |
| PILON_KLF1_TARGETS_DN [1972] | 285 | $1.37\ e^{-55}$ |
| GRAESSMANN_APOPTOSIS_BY_DOXORUBICIN_DN [1781] | 261 | $5.86\ e^{-52}$ |

*max. distance to TSS: 10,000 bp*

B

| Gene Set Name [# Genes (K)] | # Genes in Overlap (k) | p-value |
|---|---|---|
| BENPORATH_**NANOG**_TARGETS [988] | 278 | $8.72\ e^{-131}$ |
| BENPORATH_**SOX2**_TARGETS [734] | 222 | $5.03\ e^{-111}$ |
| BENPORATH_**OCT4**_TARGETS [290] | 123 | $2.26\ e^{-81}$ |
| BENPORATH_NOS_TARGETS [179] | 99 | $4.25\ e^{-80}$ |
| BENPORATH_ES_1 [379] | 113 | $8.60\ e^{-56}$ |

*max. distance to TSS: 10,000 bp*

C

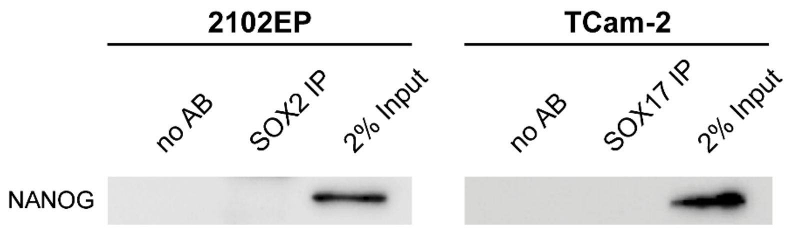


D

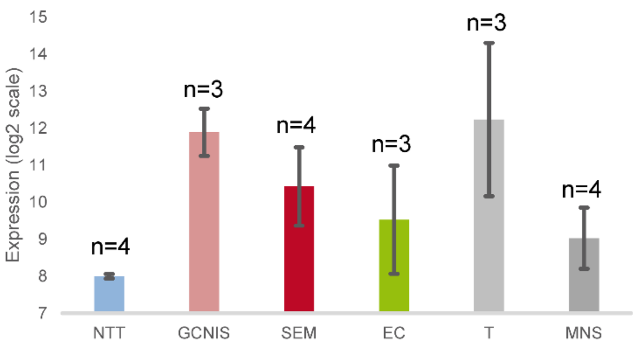


E

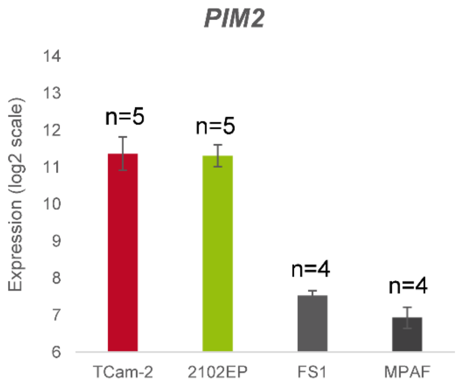


**Supplementary Figure 2**: (A) Gene set enrichment analysis (GSEA) of SOX17 ChIP-seq target genes (SOX17 signal is within a distance +/- 10,000 bp from TSS) for curated gene sets 16,17. (B) Gene set enrichment analysis (GSEA) of SOX2 ChIP-seq target genes (SOX2 signal is within a distance +/- 10,000 bp from TSS) for curated gene sets 16,17. (C) Co-immunoprecipitation of SOX17 (in TCam-2 cells) and SOX2 (in 2102EP cells) with NANOG. 2% input and no-antibody IP served as positive and negative controls, respectively. (D) Relative expression of OTX2 in normal testis tissues (NTT), GCNIS tissues, seminoma tissues (SEM), EC tissues, teratoma tissues (T) and mixed non-seminoma tissues (MNS) as determined by microarray analysis. The number of samples for each tissue type is indicated above each bar. Error bars indicate standard deviation from the mean. (E) Relative expression of PIM2 in TCam-2 and 2102EP cells, as well as a Sertoli cell line (FS1) and human adult fibroblasts (MPAF) as determined by microarray analysis. The number of samples for each tissue type is indicated above each bar. Error bars indicate standard deviation from the mean.

### Supp. Figure 3

A

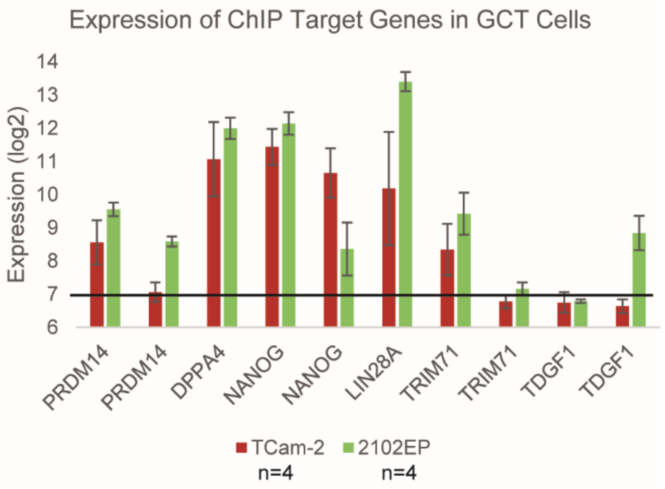


**Supplementary Figure 3**: (A) Relative expression of SOX17 and SOX2 ChIP-seq target genes in TCam-2 and 2102EP cells as determined by microarray analysis. The number of samples for each cell type is indicated below. Error bars indicate standard deviation from the mean. Different bars for the same gene indicate different probes on the DNA microarray chip.

Supp. Figure 4

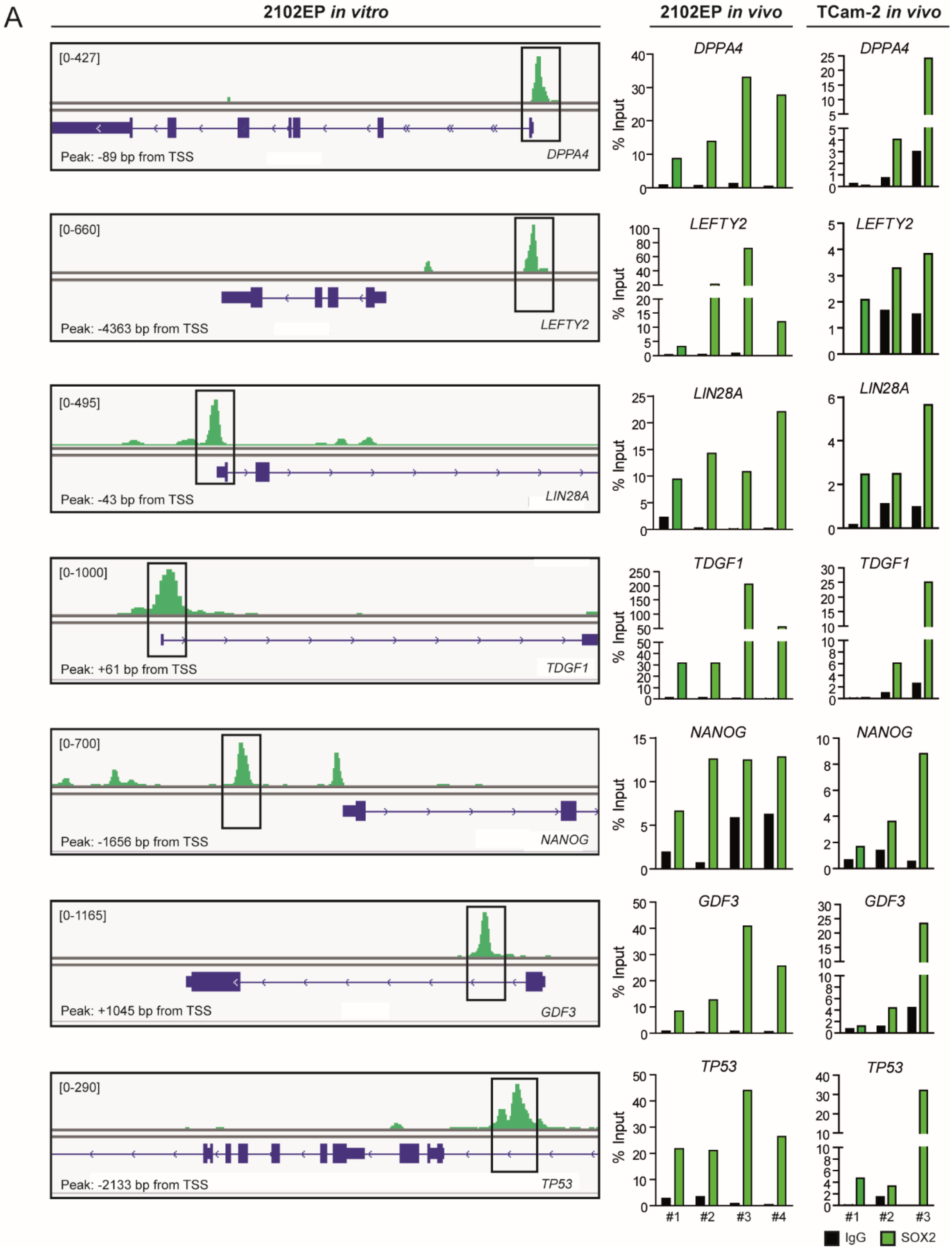

Supp. Figure 4

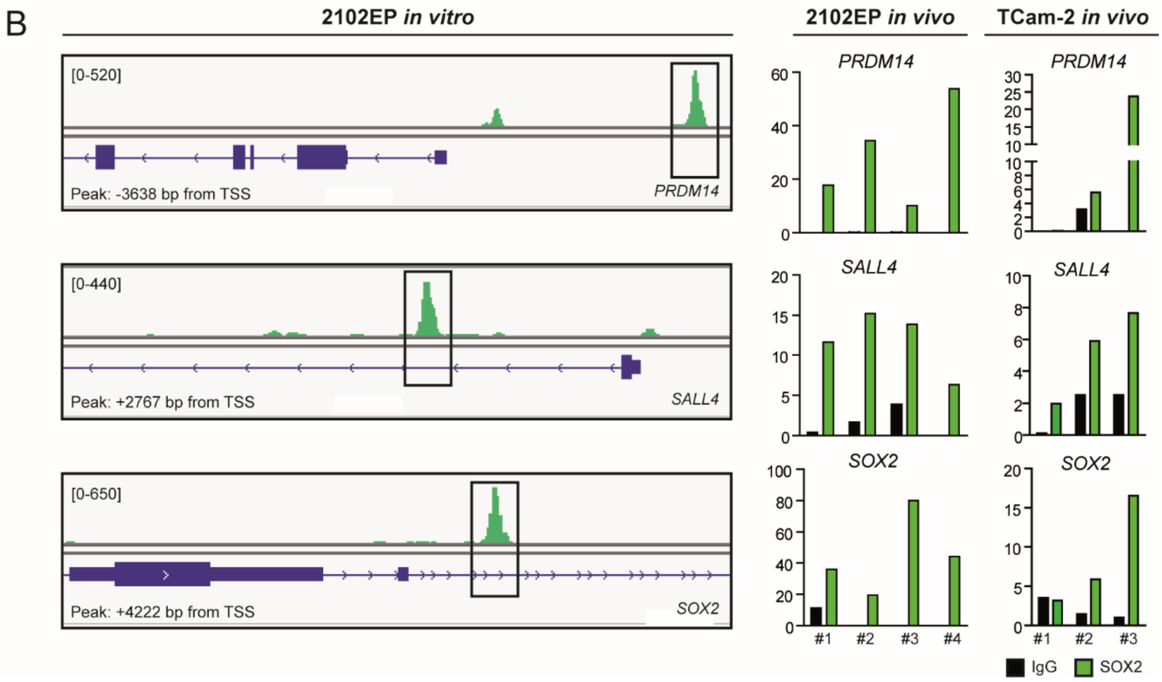


**Supplementary Figure 4:** (A-B) Left: SOX2 DNA binding profile in 2102EP cells upstream of selected promoter regions. Right: SOX2 ChIP-qPCR of 2102EP (n=4) and TCam-2 (n=3) tumour xenografts compared to IgG ChIP as negative control. Sample numbers are indicated below (#1-#4 or #1-#3). Primers are flanking the SOX2 binding sites represented in black boxes on the left. Distance of binding sites to TSS are indicated in lower left corner. Scales (upper left corner) were adjusted to allow for best visualization.

Supp. Figure 5

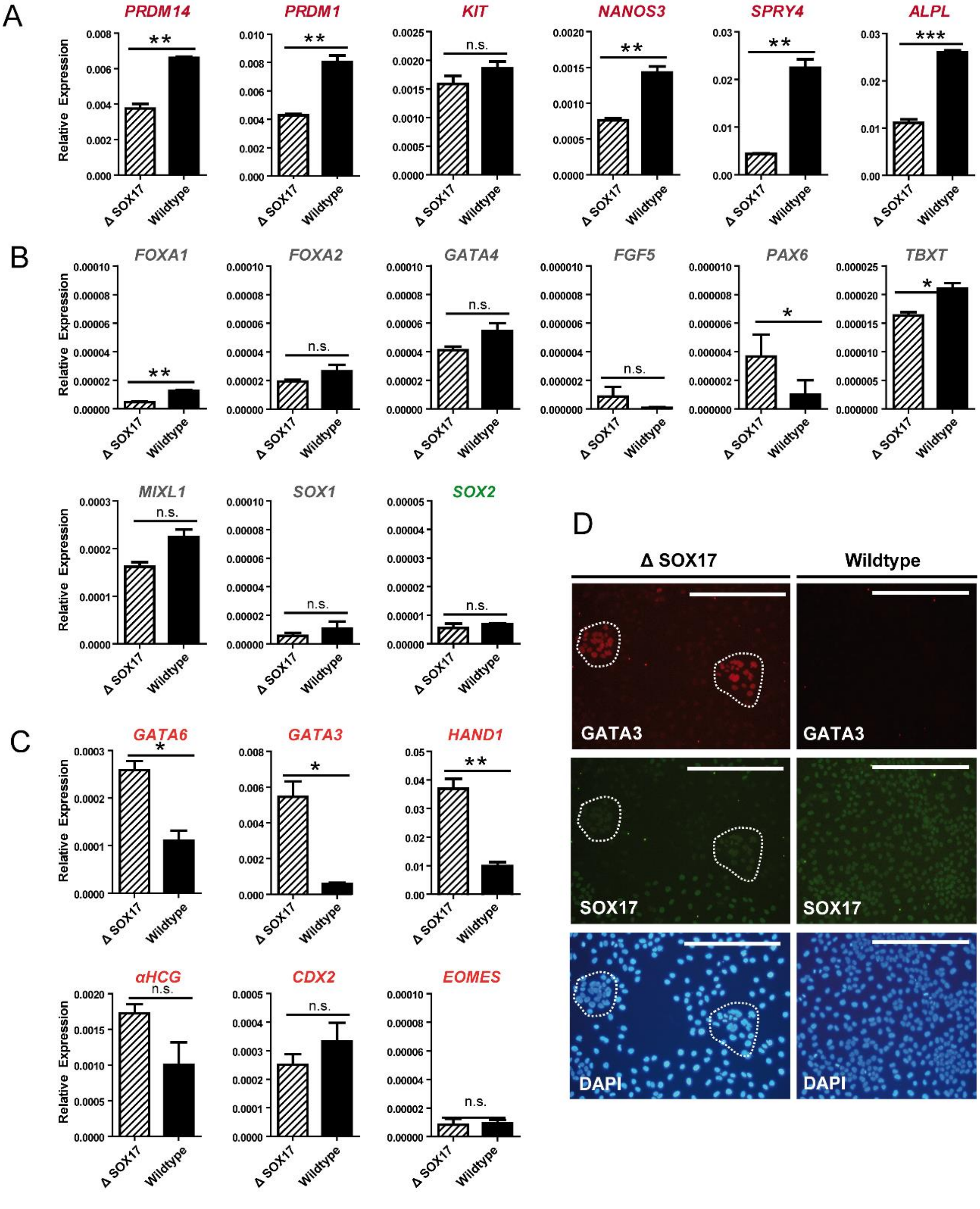


**Supplementary Figure 5**: qRT-PCR of selected germ cell markers (red labels) (A), somatic differentiation markers (grey labels) (B) and extra-embryonic differentiation markers (orange labels) (C) in TCam-2ΔSOX17 cells and TCam-2 cells transiently transfected with a GFP-coding plasmid (wildtype) as control, 72 h following transfection. Expression was normalized to GAPDH as housekeeper. Significance was calculated using two-tailed students t-test (* p ≤ 0.05; ** p ≤ 0.01; *** p ≤ 0.001). Error bars indicate standard deviation from the mean. (D) Immunofluorescence staining of GATA3 and SOX17 protein levels in TCam-2ΔSOX17 cells and TCam-2 wildtype cells. Cell nuclei were stained with DAPI. Scale bar = 250 µm.

**Supplementary Table 1:** List of primers used for ChIP-qPCR analysis

| Target Gene | Forward (5'->3') | Reverse (5'->3') | Reference |
|---|---|---|---|
| ***DPPA4*** | ACCCAGACAAAAGTCACCCC | AAGTCTCCTCCCACTTCCTG | [46] |
| ***GDF3*** | TGCTGTAAAACTAACCAAGGCTC | ACAGTAGATCCTCCAACCCC | - |
| ***LEFTY2*** | TCTCCACTCAGACCCTCAGA | GGCAGCCTGAAGAGTTTTGT | [46] |
| ***LIN28A*** | GGGTTGGGTCATTGTCTTTTAG | AAAGGGTTGGTTCGGAGAAG | [47] |
| ***NANOG*** | CAGCCCCTTCCCTTTGGTC | GGGCAGGTACCAGAAGCTTT | - |
| ***POU5F1*** | GACAATCCCGGTCCCCAGAG | GTGTGAGGGGATTGGGACTG | - |
| ***PRDM14*** | AGCAAGCCAAGCATAGGGAA | CCTAGACTGAGGCTCGTTACT | - |
| ***SALL4*** | TCCAGTTTAGCACAAAGGGCA | GGAGGAAAATGACGTCTGGC | - |
| ***SOX2*** | TGAGCACATTCGCCAGTTCT | GACTGCCTCAAATTCCGGGA | - |
| ***TDGF1*** | ATTAGAGGCCCCAGCATTCC | GCCATGTCTGCACTACAATGAC | - |
| ***TP53*** | CTCCTTCCCGTGCAGACTTT | GATAGATCACTGCCCTGCCC | - |

**Supplementary Table 2: Quality statistics of ChIP-seq analysis**

| Sample | Total Sequences (Pre-Alignment) | Total Sequences (Post-Alignment) | Unique Positions | Average Tags per Position |
|---|---|---|---|---|
| 2% Input 2102EP #1 | 27,988,206 | 24,279,589 | 22,014,628 | 1.103 |
| SOX2 2102EP #1 | 14,926,356 | 13,866,883 | 13,155,736 | 1.054 |
| 2% Input 2102EP #2 | 26,596,105 | 26,078,227 | 22,510,002 | 1.159 |
| SOX2 2102EP #2 | 35,056,570 | 31,556,291 | 28,740,374 | 1.098 |
| 2% Input 2102EP #3 | 30,945,953 | 28,460,654 | 25,100,596 | 1.134 |
| SOX2 2102EP #3 | 43,072,201 | 38,931,440 | 35,326,096 | 1.102 |
| 2% Input TCam-2 #1 | 22,713,334 | 20,752,632 | 18,889,437 | 1.099 |
| SOX17 TCam-2 #1 | 42,207,183 | 38,177,802 | 35,030,469 | 1.090 |
| 2% Input TCam-2 #2 | 26,489,150 | 24,343,239 | 22,518,757 | 1.081 |
| SOX17 TCam-2 #2 | 32,023,406 | 29,027,316 | 26,898,376 | 1.079 |
| 2% Input TCam-2 #3 | 24,635,788 | 22,504,677 | 21,579,488 | 1.043 |
| SOX17 TCam-2 #3 | 26,590,018 | 24,124,548 | 22,342,840 | 1.080 |

**Supplementary Table 3:** List of primers used for qRT-PCR analysis

| Target Gene | Forward (5'->3') | Reverse (5'->3') |
|---|---|---|
| *ALPL* | AACATCAGGGACATTGACGTG | GTATCTCGGTTTGAAGCTCTTCC |
| *CDX2* | TTCCCATCTGGCTTTTTCTG | AGAGAAGAGCTGGGGAGGAG |

| | | |
|---|---|---|
| *EOMES* | CACATTGTAGTGGGCAGTGG | CGCCACCAAACTGAGATGAT |
| *FGF5* | ATTTGCTGTGTCTCAGGGGAT | CTGTGAACTTGGCACTTGCAT |
| *FOXA1* | GCCTGAGTTCATGTTGCTGA | CTGTGAAGATGGAAGGGCAT |
| *FOXA2* | TACGTGTTCATGCCGTTCAT | CGACTGGAGCAGCTACTATGC |
| *GAPDH* | TGGTATCGTGGAAGGACTCATGAC | ATGCCAGTGAGCTTCCCGTTCAGC |
| *GATA3* | TCTGACCGAGCAGGTCGTA | CCTCGGGTCACCTGGGTAG |
| *GATA4* | ACACCCCAATCTCGATATGTTTG | GTTGCACAGATAGTGACCCGT |
| *GATA6* | AGGGCTCGGTGAGTCCAAT | CGCTGCTGGTGAATAAAAAGGA |
| *HAND1* | AAGGCTCAGGACCCAGAAG | CGGTGCGTCCTTTAATCCT |
| *KIT* | TCATGGTCGGATCACAAAGA | AGGGGCTGCTTCCTAAAGAG |
| *LIN28A* | TGTAAGTGGTTCAACGTGCG | TGTAAGTGGTTCAACGTGCG |
| *LIN28B* | GGAGCCCCTGTTTAGGAAGT | GCACTTCTTTGGCTGAGGAG |
| *MIXL1* | CTGTGCTCCTGGAACTGAAACGAAAT | ACCTTGGGAGCTAGAGTCAGAGATG |
| *NANOG* | GATTTGTGGGCCTGAAGAAA | AAGTGGGTTGTTTGCCTTTG |
| *NANOS3* | ACAAGGCGAAGACACAGGAC | AGGTGGACATGGAGGGAGA |
| *PAX6* | ACCCATTATCCAGATGTGTTTGCCCGAG | CTGCCGCCTATGCCCAGCTTCACCAT |
| *POU5F1* | CGAAAGAGAAAGCGAACCAG | GCCGGTTACAGAACCACACT |
| *PRDM1* | GGGTGCAGCCTTTATGAGTC | CCTTGTTCATGCCCTGAGAT |
| *PRDM14* | TCCACACAGGGGGTGTACTT | GAGCCTTCAGGTCACAGAGC |
| *SOX1* | ATGCACCGCTACGACATGG | CTCATGTAGCCCTGCGAGTTG |
| *SOX17* | GATGCGGGATACGCCAGTGAC | GCTCTGCCTCCTCCACGAAG |
| *SOX2* | ATGCACCGCTACGACGTGA | CTTTTGCACCCCTCCCATT |
| *SPRY4* | TCTGACCAACGGCTCTTAGAC | GTGCCATAGTTGACCAGAGT |
| *TBXT* | ATGGAGGAACCCGGAGAC | ATGAGGATTTGCAGGTGGAC |
| *TFAP2C* | CCCACTGAGGTCTTCTGCTC | AGATCACATGAGCGGCTTT |
| *αHCG* | GTGCAGGATTGCCCAGAAT | CTGAGGTGACGTTCTTTTGGA |

**Supplementary Table 4:** List of antibodies used in this study

| Antibody | Dilution | Company | # Number |
|---|---|---|---|
| GATA3 | IF: 1:200 | Santa Cruz | sc268 |
| IgG (Goat) | IP: 10 µg | Santa Cruz | sc2028 |
| IgG (Rabbit) | IP: 10 µg<br>ChIP: 5 µg | Cell Signaling | 2729 |
| LIN28A | WB: 1:1000 | R&D | AF3757 |

| | | | |
|---|---|---|---|
| NANOG | WB: 1:500 | Santa Cruz | sc134218 |
| OCT4 | WB: 1:500<br>IF: 1:100 | Santa Cruz | sc-5279 |
| SOX17 | WB: 1:1000<br>ChIP: 5 µg<br>IF: 1:25<br>IP: 10 µg | R&D | AF1924 |
| SOX2 | ChIP: 5 µg<br>IP: 10 µg | Abcam | ab59776 |
| SOX2 | WB: 1:500 | R&D | MAB2018 |
| TFAP2C | WB: 1:500<br>IF: 1:200 | Santa Cruz | sc-8977 |
| β-ACTIN | WB: 1:50000 | Sigma Aldrich | a5441 |

**Supplementary Data 1: (A)** SOX17 ChIP-seq peaks found in at least one of three replicates in TCam-2 cells, determined by HOMER (13). **(B)** SOX17 ChIP-seq peaks commonly found in all three replicates in TCam-2 cells, determined by HOMER (13). **(C)** SOX2 ChIP-seq peaks found in at least one of three replicates in 2102EP cells, determined by HOMER. **(D)** SOX2 ChIP-seq peaks commonly found in all three replicates in 2102EP cells, determined by HOMER.

**Supplementary Data 2: (A)** Top 20 Enriched Binding Motifs in SOX17-bound Regions (Peaks present in all three replicates). **(B)** Top 20 Enriched Binding Motifs in SOX2-bound Regions (Peaks present in all three replicates). **(C)** Top 20 Enriched Binding Motifs in SOX2- and SOX17- bound Regions (Peaks present in all three replicates of both SOX17 and SOX2 ChIP seq analyses).

**Supplementary Data 3: (A)** Genes regulated by SOX17 in mesoderm-, endoderm- and mesendoderm-differentiated ESCs (peaks +/- 10,000 bp from TSS) and their overlap with genes regulated by SOX17 in TCam-2 cells with a peak score > 150 and a gene expression > 7. Annotation of peaks was performed using HOMER. **(B)** Genes regulated by SOX2 in human ESCs (peaks +/- 10,000 bp from TSS) and their overlap with genes regulated by SOX2 in 2102EP cells with a peak score > 200 and a gene expression > 7. Annotation of peaks was performed using HOMER.
